# Targeting human γδ T cells as a potent and safe alternative to pan-T cells bispecific cell engagers

**DOI:** 10.1101/2023.07.10.548307

**Authors:** Lola Boutin, Clément Barjon, Laura Lafrance, Eric Senechal, Dorothée Bourges, Emmanuelle Vigne, Emmanuel Scotet

## Abstract

Over the past decade, an increasing number of immunotherapies aiming to improve the ability of the immune system to effectively eradicate tumor cells have been developed. Among them, targeting effector T cell subsets of the immune system with bispecific antibodies, called T Cell Engagers (TCEs), represents an attractive strategy. TCEs are designed to specifically direct cytotoxic T cells towards tumor cells, thereby inducing a strong activation leading to the lysis of tumor cells. New strategies for targeting specific T-cell subsets are currently being explored. In this study, we investigated the activity of different TCEs on both conventional alpha beta (αβ) T cells and unconventional gamma delta (γδ) T cells. We generated TCE molecules based on camelid single-domain antibodies (VHHs) that target the tumor-associated antigen CEACAM5 (CEA), together with particular T-cell receptor chains (TCRs) or a CD3 domain. The *in vitro* biological activity of the TCEs against the colon carcinoma cell line LS174T was measured using fresh and cultured human Vγ9Vδ2 and αβ T cells. We showed that Vγ9Vδ2 T cells display stronger antitumor activity *in vitro* than αβ T cells when activated with a CD3xCEA TCE. Furthermore, restricting T cell activation to Vγ9Vδ2 T cells limits the production of pro-tumor factors and pro-inflammatory cytokines, which are often associated with toxicity in patients. Taken together, these results suggest that Vγ9Vδ2γδ T cell-specific TCEs may represent safe, novel, specific, and effective molecules for improving antitumor immunotherapies.

## Introduction

The persistence and increase in cancer deaths worldwide partly reflects the failure of conventional treatments such as chemotherapy, radiation, and surgery. Over the last decade, redirecting immune cells against tumor cells using bispecific antibodies has been a promising and attractive strategy among antitumor immunotherapies. T Cell Engagers (TCEs) are bispecific molecules designed to bind both T cells via T cell receptors and infected or transformed target cells via specifically expressed antigens^1^. Simultaneous binding to the two types of cells induces the formation of an active immune synapse and leads to the death of the target cell without the need for costimulatory signals^2–4^ while bypassing tumor escape mechanisms such as major histocompatibility complex (MHC) downregulation. These molecules can be primarily classified according to the presence or absence of an Fc region, number of binding sites, and overall geometry, which together govern their pharmacokinetic and pharmacodynamic properties^5^. Their antigen-binding sites may arise from conventional two-chain immunoglobulins or from single-domain antibodies, also known as VHHs, of camelid origin^6–8^. VHHs are modular building blocks characterized by their small size (15 kDa) and excellent biophysical properties, which enable their easy assembly into multispecific functional molecules. They are well suited for therapy, as proven by the recent approval in Japan of Ozoralizumab, a trivalent anti-TNFα Nanobody^Ⓡ^ compound, for the treatment of rheumatoid arthritis^9^.

Initial studies highlighted a therapeutic benefit of TCEs in the treatment of hematological malignancies, leading to the approval of Blinatumomab (CD3xCD19) or Mosunetuzumab (CD3xCD20) in the treatment of acute lymphoblastic B cell leukemia and follicular lymphoma respectively^10, 11^. More recently, clinical trials using TCEs targeting the prostate-specific membrane antigen (PSMA) or human epidermal growth factor receptor 2 (HER2) have reported promising results in solid tumors^12, 13^. A recent study has further demonstrated the benefits of combining TCE therapy with immune checkpoint blockade (ICB) and a 4-1BB agonist in T-cell cold solid tumors, where ICB monotherapy did not show any therapeutical benefit^14–16^. The majority of current TCEs target T cells via the CD3 signaling complex, thus inducing strong polyclonal T cell activation. TCEs were initially designed to trigger minimal side effects by promoting tumor site-specific T cell recruitment, infiltration, and activation. However, stage 3 (or higher) deleterious effects in patients, such as Cytokine Release Syndrome (CRS), neurotoxicity, and neutropenia, are frequently associated with treatment^17^. In addition, the use of such molecules might also increase the risk of activating pro-tumor, inhibitory, or regulatory T cells (Tregs), which could reduce the efficacy of these immunotherapies^18, 19^.

TCEs that specifically activate selected T cell subsets such as γδ T cells have been proposed to address these important issues^20–24^. In humans, γδ T cells are distributed into four major subsets identified according to the expression of the T Cell Receptor (TCR) δ chain variable segment: Vδ1^+^, Vδ2^+^, Vδ3^+^, and Vδ5^+^ populations. The major peripheral blood subset in healthy human adults is highly conserved, devoid of alloreactivity, and expresses a TCR composed of a Vδ2 chain that is almost exclusively associated with a TCR Vγ9 chain, whereas Vδ1^+^ and Vδ3^+^ T subsets are most frequently found in tissues^25^. A compelling set of studies has shown that Vγ9Vδ2 T cells can directly kill tumor cells and express pro-inflammatory cytokines involved in the clearance of tumor cells, supporting the development of immunotherapies engaging this lymphocyte subset^26–30^.

In this study, we challenged conventional αβ T cells and unconventional γδ T cells using TCEs directed against the tumor antigen CEACAM5 (CEA) in an *in vitro* colorectal cancer model. Our results show that the particular biology of γδ T cells translates into a differential TCE-mediated antitumor response over conventional αβ T cells. Moreover, our results suggest that specific targeting of Vγ9Vδ2 T cells by TCE would promote an enhanced therapeutic safety by reducing cytokine release syndrome.

## Materials and methods

### T cell Engager molecules

VHH sequences directed against TCRγδ, TCRαβ, and CD3 were retrieved from patent applications WO2015/156673 (clones #5C8; #6H1), WO2016/180969 (clone #56G05) and WO2016/180982 (clone #117G03), respectively. VHH sequences directed against CEACAM5 and FMDV have been previously published^31, 32^. DNA sequences for VHHs were synthesized by GeneArt (Life Technologies) as string DNA fragments and cloned into a eukaryotic expression vector fused with human CH1 or CL domains. FreeStyle™ HEK293-F cells (Gibco) were co-transfected with two plasmids, each encoding one of the Fab chains. After seven days, the culture media were harvested and filtered before purification of the constructs on nickel-affinity columns.

### Reagents

AF647-conjugated anti-CD107a, AF647-conjugated anti-perforin, PE-conjugated anti-PD1 and FITC-conjugated anti-Vγ9 mAbs were purchased from BioLegend. BV510-conjugated anti-CD4 and BV421-conjugated anti-CD8 mAbs were purchased from BD Bioscience. FITC-conjugated anti Vδ2 and PE-conjugated anti CD69 mAbs were purchased from Beckman Coulter. APC-conjugated anti-Vδ1 mAb (REA) was purchased from Miltenyi. AF568-conjugated phalloidin was purchased from Life Technologies.

### Cells

LS174T (human colon carcinoma), LS174T_shGFP (LS174T cell line stably transduced with lentiviral vector FG12.34 expressing short half-life variant green fluorescent protein), HEK293 (human embryonic kidney 293), and HEK EBNA_CEA^+^ (HEK293 cell line stably expressing Epstein-Barr virus nuclear antigen-1 and CEA) cells were grown in low-glucose (1g/L) DMEM-GlutaMax medium (Life Technology) supplemented with 10% heat-inactivated fetal calf serum (FCS) in a humidified chamber at 37°C and 5% CO_2_. HEK293 (CRL-1573) and LS174T (CL-188) cells were purchased from the American Type Culture Collection (ATCC). FreeStyle™ HEK293-F cells were cultured in Freestyle™ 293 expression medium (Gibco).

### Isolation and amplification of human T cells

Human peripheral blood mononuclear cells (PBMCs) were isolated from blood of healthy donors (informed consent) obtained from Etablissement Français du Sang (Nantes, France). PBMCs were collected by Ficoll gradient centrifugation and resuspended in RPMI-1640 medium supplemented with 5% heat-inactivated FCS. T cells were enriched from PBMCs by 30 min-incubation at 37^◦^C in a horizontal tissue-culture flask (up to 7.10^6^ cells/mL) to promote the adhesion of monocytes, macrophages, and B cells. T cell-enriched supernatants were obtained by centrifugation and used directly or were frozen in heat-inactivated FCS plus 10% DMSO.

Human Vγ9Vδ2 T cells were specifically expanded from fresh PBMCs *ex vivo* using 5 µM zoledronic acid (Sigma Aldrich) or 3 µM BrHPP (bromohydrin pyrophosphate, kindly provided by Innate Pharma) in RPMI 1640 culture medium supplemented with 10% heat-inactivated FCS, 2 mM L-glutamine, 100 µg/mL streptomycin and 100IU/mL recombinant human IL-2 (rhIL-2) (Proleukin®, Novartis). After 4 days of culture, T lymphocytes were supplemented with rhIL-2 (300 IU/mL). After 3 weeks (resting state time), the purity of Vδ2^+^ T cells was checked using flow cytometry (purity range = 85-95%). Non-specific amplifications were performed using PHA-feeders: Phytohemagglutinin-L (PHA-L, Sigma-Aldrich) and 35 Gy-irradiated allogeneic feeder cells composed of human PBMC and Epstein-Barr virus-transformed B-lymphoblastoid cell lines^33^. Expanded Vγ9Vδ2 T cells were maintained in RPMI 1640 culture medium supplemented with 10% heat-inactivated FCS, 2 mM L-glutamine, 100 µg/mL streptomycin and 300 IU/mL rhIL-2.

Human αβ T cells were expanded from fresh PBMCs by incubation with ImmunoCult® human CD3/CD28 Activator (StemCell) in RPMI 1640 culture medium supplemented with 10% heat-inactivated FCS, 2 mM L-glutamine, 100µg/mL streptomycin, and 100 IU/mL rhIL-2. After 12 days of expansion, the purity of the amplified αβ T cells was checked using flow cytometry (purity range = 85-95%).

### T cell activation assays

For CD107a surface mobilization assays, target cells were co-cultured for 4 h with amplified T cells (E/T ratio 1:1) in the presence of TCE (concentration range:10 fM-1 nM) in culture medium containing 5 µM Monensin (Sigma Aldrich) and anti-human CD107a mAb. For CD69 expression, LS174T cells were co-cultured with fresh T cell-enriched PBMCs for 24 h (E/T ratio 10:1) in the presence of TCE (concentration range: 10 fM-1 nM). Extracellular marker staining was performed after co-culture by incubating with the antibody mixture for 20 min at 4°C. Flow cytometry data were acquired using the Accuri C6 or Canto II cytometer (BD Biosciences) and analyzed using FlowJo software (Treestar).

### Cytotoxicity assays

LS174T cells were incubated with ^51^Cr (75µCi/1.10^6^ cells) for 1 h, washed, and co-cultured with *ex vivo* expanded T cells (E/T ratio 10:1) for 4 h in the presence of TCE (concentration range:10 fM-1 nM). ^51^Cr-release in the supernatant was measured using a MicroBeta counter (Perkin Elmer). The percentage of target cell lysis was calculated as follows: ((experimental release - spontaneous release)/ (maximum release - spontaneous release)) × 100. Spontaneous and maximum release values were determined by adding either medium or Triton X-100 to ^51^Cr-labeled target cells without T cells.

### Imaging of the immunological synapse

5.10^5^ LS174T-GFP cells were plated in Ibidi µ-Slide 8 wells (Ibidi GmbH) coated with CellTak adhesion matrix (Corning) overnight at 37°C. TCE (1 nM) and *ex vivo* expanded T cells (E/T ratio 10:1) were then added for 30 min. The cells were fixed in 4% paraformaldehyde for 10 min at room temperature (RT) and permeabilized in PBS containing 0.1% BSA and 0.1% saponin for 20 min at RT. Non-specific binding sites were saturated with PBS containing 10% FCS and 0.1% saponin for 20 min at RT. The cells were then labeled with anti-human perforin and phalloidin for 1 h at RT in PBS containing 0.1% BSA and 0.1% saponin. Imaging was performed using a Nikon A1 confocal microscope [60X oil immersion objective, N.A 1.4]. The distance between the perforin granules and the contact area between the tumor and T cells was measured using NIS analysis software (Nikon).

### Cytokine/granzyme B bead-based assays

Supernatants from experiments of enriched T cell activation were collected after 24 h of co-culture with LS174T cells (E/T ratio 10:1) in the presence or absence of TCE (concentration specified for each experiment). Concentrations of released IL-2, IL-10, IL-6, TGF-β1, IFN-γ, TNF-α, and granzyme B were measured using bead-based flow cytometry (LegendPlex, BioLegend) according to the manufacturer’s protocol. Data were acquired with the Accuri C6 cytometer or the Canto HTS cytometer (BD Biosciences) and analyzed using the LegendPlex software (BioLegend).

## Results

### Human αβ and γδ T cells react differently following CD3 engagement

Various studies have shown that human γδ and αβ T cells differ on their development, phenotype, and effector functions^25^. To investigate the underlying functional features, we compared the effects of a soluble anti-CD3ε mAb on the activation of *ex vivo*-expanded human Vγ9Vδ2 and αβ T cells. As shown in **Fig. 1A** and **Fig. 1B**, anti-CD3ε mAb (#UCHT1) induced higher levels of activation (monitored by CD107a cell surface expression) in Vγ9Vδ2 T cells than in αβT cells, at both low and high mAb concentrations. Furthermore, UCHT1-mediated activation of different γδ and αβ T cell subsets enriched from fresh peripheral blood (Vδ2^+^/Vδ1^+^γδ T cells, CD8^+^/CD4^+^ αβ T cells) demonstrated a stronger reactivity of Vγ9Vδ2 T over Vδ1^+^ and αβ T cells as revealed by the comparative analysis of CD69 expression levels at cell surface (**Fig. 1C** and **Fig. 1D**). These results indicate that Vγ9Vδ2 and αβ T cell subsets react differently upon CD3ε stimulation, the former unconventional T cell subset requiring significantly lower amounts of anti-CD3ε mAb to reach strong activation levels.

**Figure 1:**
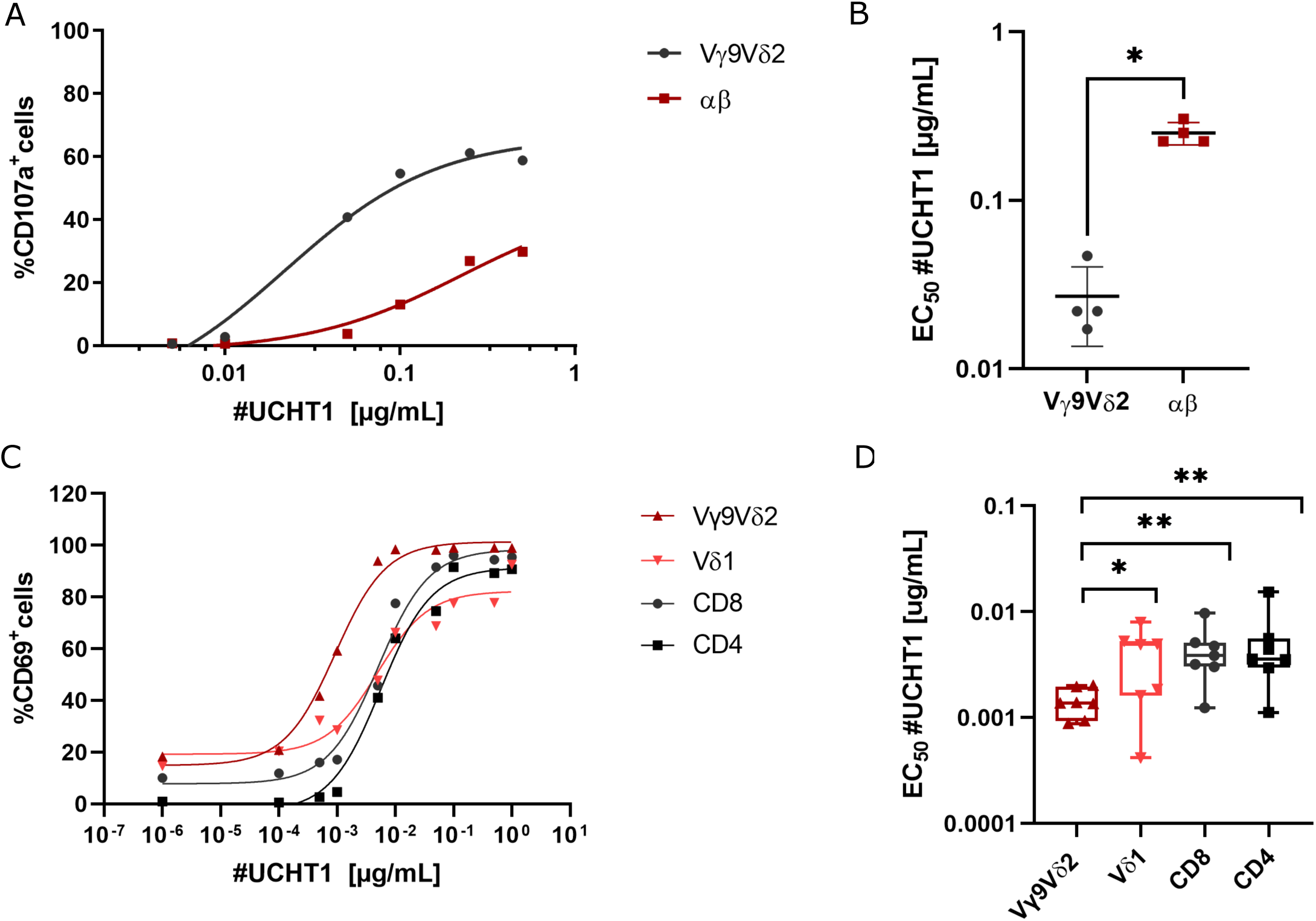
Anti-CD3 mAb induces a stronger activation of human γδ T cells, as compared to αβ T cells. (**A**) Analysis by flow cytometry of surface expression of CD107a *ex vivo*-expanded human Vγ9Vδ2 and αβ T cells after a 4 h-activation with soluble anti-CD3 mAb (#UCHT1). (**B**) Calculated activation potential (EC_50_) from experiments shown in (A) panel; n=4 unpaired donors. Unpaired Mann-Whitney (non-parametric) test used to assess the significance of differences between calculated EC_50_ for T cell subsets. (**C**) Analysis by flow cytometry of CD69 surface expression on selected T cell populations from fresh T-cell enriched PBMCs after a 24 h-activation with soluble anti-CD3 mAb (#UCHT1). **(D)** Calculated activation potential (EC_50_) from experiments shown in (C) panel; n=7 donors. Statistical analysis was performed using one-way ANOVA tests on log-transformed data with cell population as fixed effect and donor as random effect, followed by Newman-Keuls tests to correct for multiplicity. (A) and (C): representative graphs from representative donors. (B) and (D) Data show box and min to max whiskers plot.

### Generation and validation of T cell engager molecules

Based on these results, TCE molecules were engineered with a VHH1 domain targeting either human CD3 or TCR (Vδ2, Vγ9 or αβTCR) subunits, and a VHH2 domain targeting CEA. CEA is overexpressed in most human colorectal, pancreatic, lung or breast carcinoma and is a validated tumor antigen for solid cancer immunotherapies^34–37^. The CEA VHH2 was swapped with an irrelevant VHH2 binding foot-and-mouth disease virus (FMDV) to serve as a negative control. VHH1 and VHH2 were combined in a bispecific Fab-*like* format (bsFab), which relies on the substitution of the VH and VL domains of a conventional Fab with two independent VHH domains^38^. HA and His tags were further appended to the C-terminus of the CH1 domain to allow optimal detection and purification of TCE molecules (**Fig. 2A**). The apparent affinities (EC_50_) of bsFabs were determined on CEA-expressing tumor cells LS174T and expanded Vγ9Vδ2 or αβ T cells (**Table 1**). CD3 bsFabs displayed close affinity values for both Vγ9Vδ2 and αβ T cells (3.9 nM *vs* 6.9 nM for CD3xCEA; 97 nM *vs* 16 nM for CD3xFMDV). Importantly, bsFabs with VHH1 targeting TCR subunits specifically bound to their associated T cell subsets albeit with contrasted affinity values (approximately 100 nM for αβTCR bsFabs *vs* nM range for Vδ2- and Vγ9-based bsFabs). Finally, all CEA bsFabs bound LS174T tumor cells with strong affinity, whereas as expected, the FMDV-based bsFabs did not bind these cells. Both the specificity and functionality of bsFabs were then assessed in a co-culture assay using HEK293 cells, either wild-type or stably expressing CEA. Expanded Vγ9Vδ2 T cells were strongly activated by Vγ9-, Vδ2-, or CD3xCEA bsFabs (**Fig. 2B**) when co-cultured with CEA-HEK293 cells. αβ T cells were activated to a lesser extent by αβTCR- or CD3xCEA bsFabs (**Fig. 2C**). For all bsFabs, the activation was dose-dependent and only detected in the presence of CEA-positive cells. These results demonstrate that all bsFabs are functionally active and have the capacity to activate the desired T cell subsets in a tumor-antigen dependent manner.

**Figure 2:**
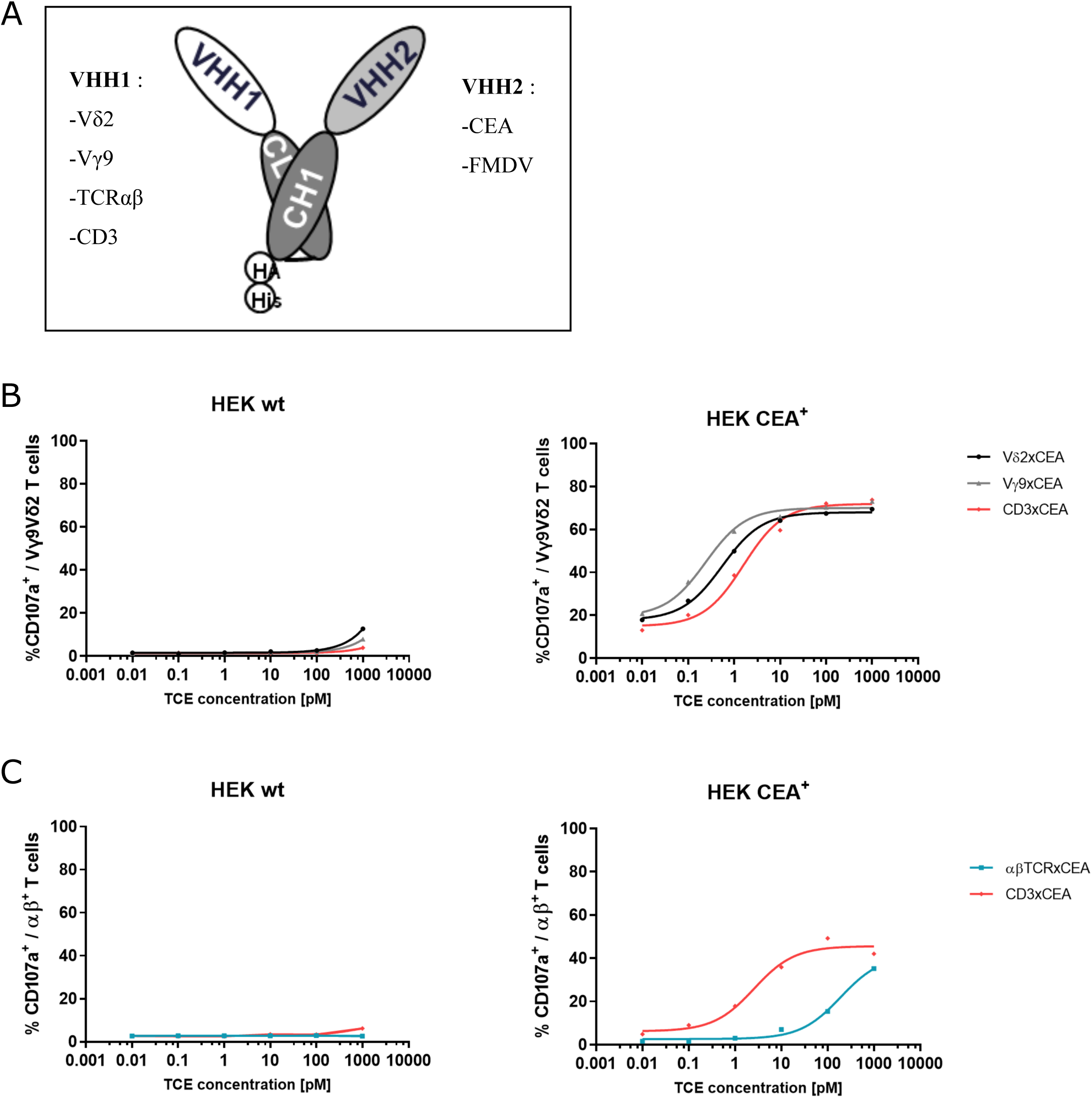
Design and validation of TCE molecules. (**A)** Schematic representation of the Fab- *like* TCE format and bispecific target design between VHH1 (effector T cell): Vδ2, Vγ9, αβ TCR or CD3, and VHH2 (target cell): CEACAM5 (tumor antigen) or FMDV (negative control) fused to human CL and CH1 chains, respectively. (**B**) Flow cytometry analysis of CD107a expression on human *ex vivo*-expanded Vγ9Vδ2 T or (**C**) αβ T cells after 4 h of co-culture with wild-type or CEACAM5-expressing HEK293 cells (E/T ratio 1:1) plus xCEA bsFabs; n=3 donors. Data from a representative donor.

**Table 1:** EC_50_ calculated from binding curves of TCEs on LS174T cell line, ex-vivo expanded Vγ9Vδ2 and αβ T cells. Values are expressed in nM

|  | *Vγ9Vδ2 T* |  | *$\alpha\beta$ T* |  | *LS174T* |  |
| --- | --- | --- | --- | --- | --- | --- |
| *VHH* | xCEA | xFMDV | xCEA | xFMDV | xCEA | xFMDV |
| Vδ2 | 1.3 | 0.2 | >1000 | >1000 | 1.3 | >1000 |
| Vγ9 | 0.6 | 4.6 | >1000 | >1000 | 1.8 | >1000 |
| TCR$\alpha\beta$ | >1000 | >1000 | 107 | 84.4 | 0.8 | >1000 |
| CD3 | 3.9 | 96.6 | 6.9 | 15.7 | 0.02 | >1000 |

### *Ex vivo*-expanded human Vγ9Vδ2 T cells are activated by bsFabs in the presence of CEA-expressing tumor cells

The human colon carcinoma cell line LS174T, which expresses endogenously CEA, was further used to test, and compare the specificity and potency of bsFabs in activating *ex vivo*-expanded Vγ9Vδ2 and αβ T cells in a dose-response assay. LS174T and T cells were co-cultured (E/T ratio: 1:1) in the presence of CEA or FMDV bsFabs. T cell activation was monitored by measuring the CD107a membrane expression. As shown in **Fig. 3A**, Vδ2-, Vγ9- and CD3xCEA bsFabs activated Vγ9Vδ2 T cells with sub pM potency (EC_50_= 0.6 ± 0.24 pM; 0.47 ± 0.17 pM; 0.56 ± 0.42 pM, respectively). In contrast, FMDV TCEs induced very low level of activation, only at the highest tested doses (∼1 nM) (**Suppl Fig. 1A**), confirming that bsFabs were active in a tumor antigen-dependent manner. In addition, Vδ2- and Vγ9xCEA bsFabs specifically activated Vγ9Vδ2 T cells with equivalent potency and efficacy and failed to activate αβ T cells. αβ T cells were exclusively activated by either αβTCR- or CD3xCEA bsFabs. However, a 50-fold higher potency was measured for CD3 bsFab over the αβTCR bsFab (EC_50_= 1.8 ± 0.12 pM *vs* 47.7 ± 2.7 pM). Moreover, the plateau of αβ T cell activation induced by CD3xCEA bsFab was much lower than that of Vγ9Vδ2 T cells (55 ±0.46 % *vs* 87 ±5,6 %).

**Figure 3:**
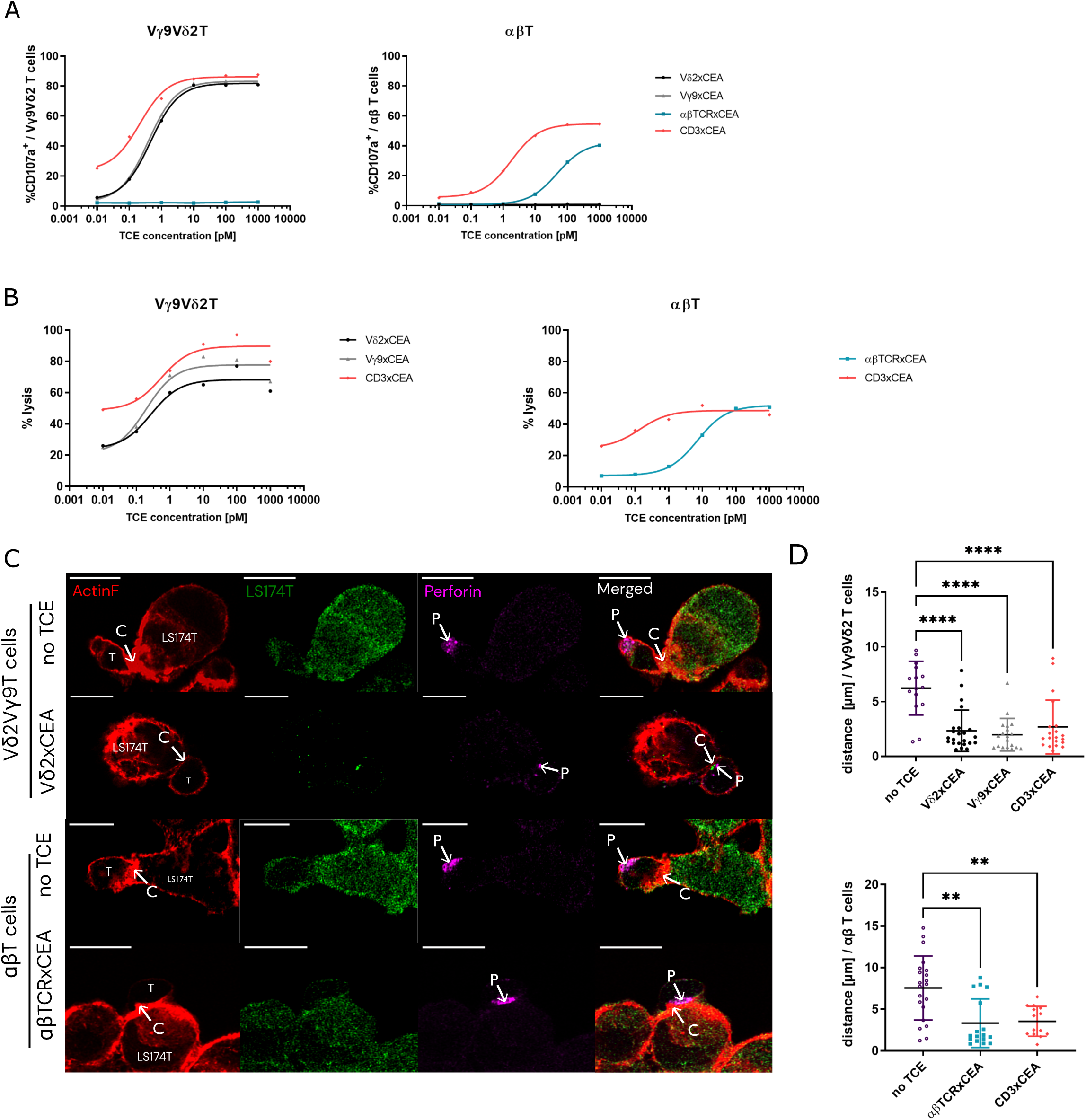
Comparison of bsFabs activation potency between *ex vivo* expanded Vγ9Vδ2 and αβ T cells. (**A**) Analysis by flow cytometry of CD107a expression on *ex vivo*-expanded human Vγ9Vδ2 or αβ T cells after 4 h of co-culture with LS174T cells (E/T ratio 1:1), plus xCEA bsFabs. n=3 donors. (**B**) LS174T cells were loaded with ^51^Cr and co-cultured for 4 h with *ex vivo*-expanded human Vγ9Vδ2 or αβ T cells (E/T ratio 10:1), plus xCEA bsFabs. ^51^Cr release was measured in culture supernatants; n=4 unpaired donors. (**C**) Confocal microscopy images of Vγ9Vδ2 and αβ T cells co-cultured with LS174T_GFP (green) for 30 min (E/T ratio 10:1) in the presence or absence of Vδ2-, or αβTCRxCEA TCEs (1nM), before staining with phalloidin-AF568 (ActinF, red) and anti-human perforin antibody-AF647 (magenta). White arrows {C=contact, P=perforin}; scale bars=10 µm. (**D**) Measurement of the distance between perforin granules and the contact point between Vγ9Vδ2 or αβ T cells and LS174T (E/T ratio 10:1) from confocal microscopy images in the presence or absence of xCEA bsFabs (1nM); n=12 to 23 images per condition. Statistical analysis was performed using one-way ANOVA test (** *P<* 0.003; **** *P<* 0.0001). (A), (B) and (C) Data presented from one representative donor. (D) Data presented are means ± SD.

Following cell expansion, the relative CD8 and CD4 T cell frequency varied substantially between donors (**Suppl Fig. 2A**). We measured separately the activation of both CD8^+^ and CD4^+^ αβ T cells upon treatment with αβTCR- and CD3xCEA bsFabs (**Suppl Fig. 2B**) and observed a dramatic difference between the maximal percentage of activated CD4^+^ αβ T and CD8^+^ αβ T cells in response to each bsFab (25.89 ± 3.7 vs 58.32 ± 5.1 % for αβTCR bsFab; 39.69± 4.2 vs 71.95 ± 3.4 % for CD3 bsFab). Of note, the CD8^+^ αβ T cell subset showed a slightly lower activation efficacy (71.95 ± 3.4 % *vs* 87.10 ± 5.6 %) and potency (0.97 ± 0.4 % *vs* 0.56 ± 0.4 pM) than Vγ9Vδ2 T cells when activated with the CD3xCEA bsFab.

The T cell-directed cytotoxicity induced by bsFabs against LS174T was next measured (**Fig. 3B).** At E/T ratio of 10:1, Vγ9Vδ2 T cells potently killed tumor cells at sub pM concentrations of Vδ2-, Vγ9- and CD3xCEA TCEs (*left panel*, EC_50_= 0.6 ± 0.24 pM *vs* 0.47 ± 0.17 pM *vs* 0.56± 0.42 pM respectively) in a tumor-antigen dependent manner (FMDV bsFab effect shown in **Supp Fig. 1B** *left*). Similarly, αβ T cells killed tumor cells in the presence of CEA bsFabs only (CEA bsFabs in **Fig. 3B** *right* and FMDV bsFab in **Supp Fig. 1B** *right*). A 26-fold difference in potency was measured between CD3- and αβTCRxCEA bsFabs (EC_50_= 0.2 ± 0.4 pM *vs* 5.2 ± 5.2 pM respectively) and the maximal killing levels by αβ T cells was much weaker as compared to Vγ9Vδ2 T cells when activated with the CD3xCEA bsFab (39.9 ±3.7 *vs* 82 ±3.8%, respectively).

To investigate the effects of bsFabs in more detail, expanded T cells were co-cultured with GFP-transduced LS174T (E/T ratio 10:1), in the presence or absence of CEA bsFabs, and stained for F-actin and perforin. This allows monitoring of the translocation of perforin granules towards the contact area established between T cells and target cells as a marker of an active immunological synapse (**Fig. 3C**). In the absence of TCE bsFabs, the intracellular pool of perforin granules (P) was located on the opposite side of the contact zone (C). By contrast, when CEA bsFabs were added to the co-culture, perforin granules migrated rapidly (within 30 min) towards the contact zone, resulting in a significant decrease in the distance between perforin granules and the immunological synapse regardless of the bsFabs and T cell subset (**Fig. 3D**).

Altogether, these results show that bsFabs mediate specific interactions between T and tumor cells in an antigen-dependent manner, resulting in their lysis via a perforin-based mechanism.

### Both CD3 and TCR-specific bsFabs strongly activate human γδ T cells in PBMCs

The stimulatory activity of bsFabs was investigated using fresh T cell enriched PBMCs isolated from the peripheral blood of healthy donors at E/T ratio of 10:1. The results showed that CEA bsFabs activate T-cell subsets in both a dose-dependent and VHH-specific manner when co-cultured with CEA-expressing cells (**Fig. 4A**). They also confirmed that the CD3xCEA bsFab activated CD4^+^ and CD8^+^ αβ T cells more than the αβTCRxCEA bsFab, whereas the stimulatory activity of Vγ9-, Vδ2-, and CD3xCEA bsFabs on Vγ9Vδ2 T cells was comparable. Importantly, the calculated EC_50_ values of the CD3xCEA TCE for each T cell subset showed an increased potency to activate Vγ9Vδ2 T cells, as compared with other T cell subsets, including CD8^+^ αβ T cells (**Fig. 4B**). Co-culture supernatants were collected 24 hours after treatment with the EC_90_ concentration of each TCE (Table 2), and the level of secreted pro- or anti-inflammatory cytokines was measured (Fig. 4D) (**Fig. 4D)**. While the CD3xCEA TCE and OKT3 mAb (CD3 control) induced a similarly strong release of cytokines, both Vγ9- and Vδ2xCEA bsFabs triggered significantly lower production of IFN-γ, TNF-α, and IL-6 (**Fig. 4C**). These cytokines were measured at a similar level after αβTCRxCEA bsFab treatment, despite the lower E/T ratio of Vγ9Vδ2 T over αβ T cells (approximatively 7 vs 0.4) in the co-cultures. Moreover, no IL-2 was detected in the co-culture supernatants when Vγ9Vδ2 T cells were activated by TCR-specific TCEs.

**Figure 4:**
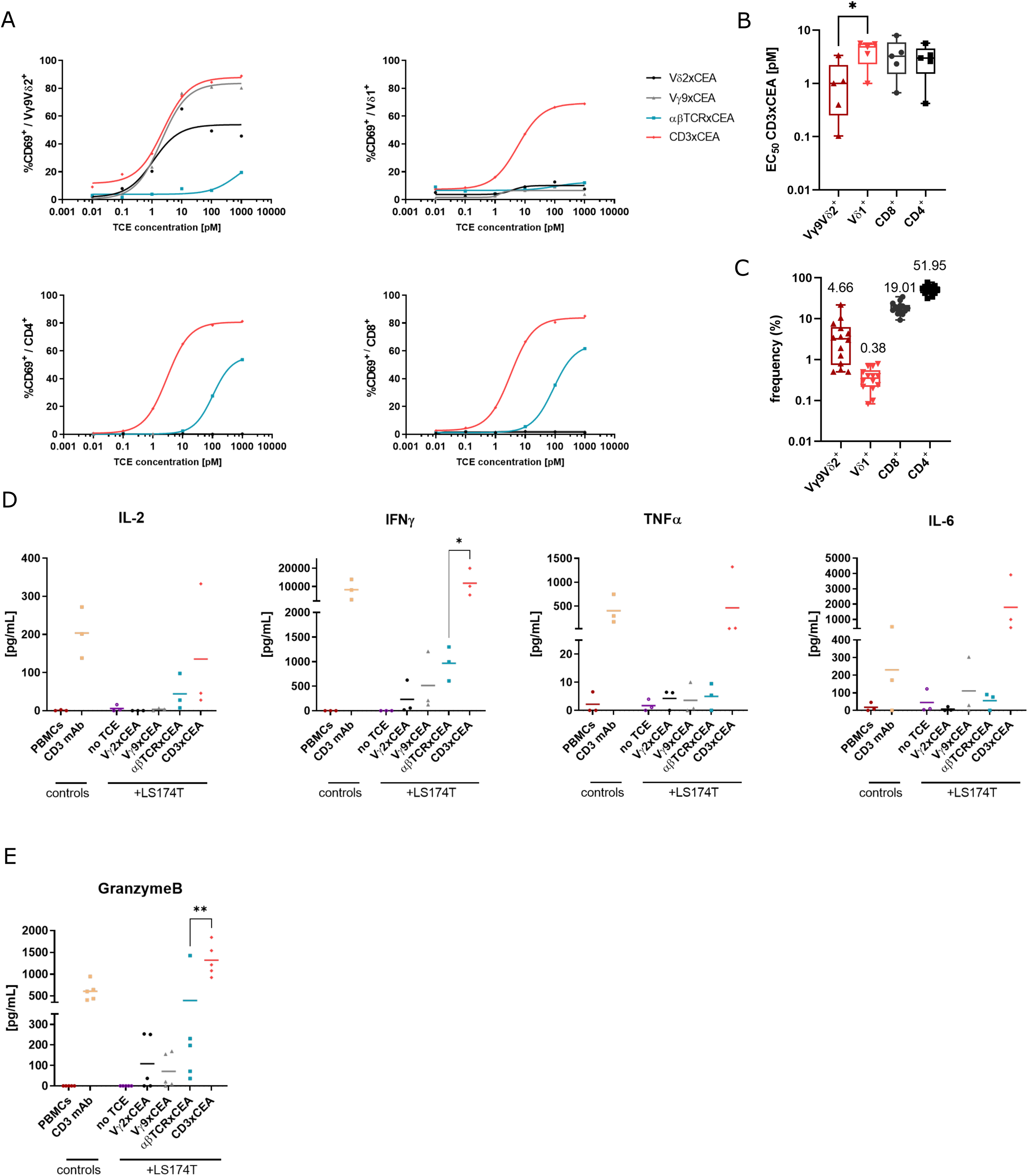
CEA bsFabs specifically activate human peripheral T cells. (**A**) Analysis by flow cytometry of CD69 surface expression on fresh PBLs (gates: Vδ2^+^, Vδ1^+^, CD4^+^ or CD8^+^ T cells) after 24 h-co-culture with LS174T target cells (E/T ratio 10:1), in the presence of indicated xCEA bsFabs; n=5 donors. The data presented from one representative donor. (**B)** Calculated activation potential (EC_50_) from experiments with CD3 bsFab (calculated from fig4.A); n=5 donors. Statistical analysis was performed using one-way ANOVA test (* *P*<0.05). (**C**) Frequencies of Vδ2^+^, Vδ1^+^, CD4^+^, and CD8^+^ T cell subsets in fresh T-cell enriched PBMCs measured by flow cytometry; n=14 donors. (**D**) IFN-γ, IL-2, TNF-α and IL-6 concentration measured in the 24 h supernatant of fresh T-cell enriched PBMCs co-cultured with LS174T cells (E/T ratio 10:1), in the presence or absence of xCEA bsFabs at their EC_90_ (calculated from fig.4A); n=3 donors. (**E)** Granzyme B release in the 24 h supernatant of fresh T-cell enriched PBMCs co-cultured with LS174T cells (E/T ratio 10:1) in the presence or absence of xCEA bsFabs (1nM); n=5 donors. (B) and (C) Data show box and min to max whiskers plot. (D) and (E) Data presented are mean - statistical analysis performed by one-way ANOVA tests, followed by two-tailed Dunnett’s tests (\**P*< 0.05; \*\**P*< 0.01).

**Table 2:**
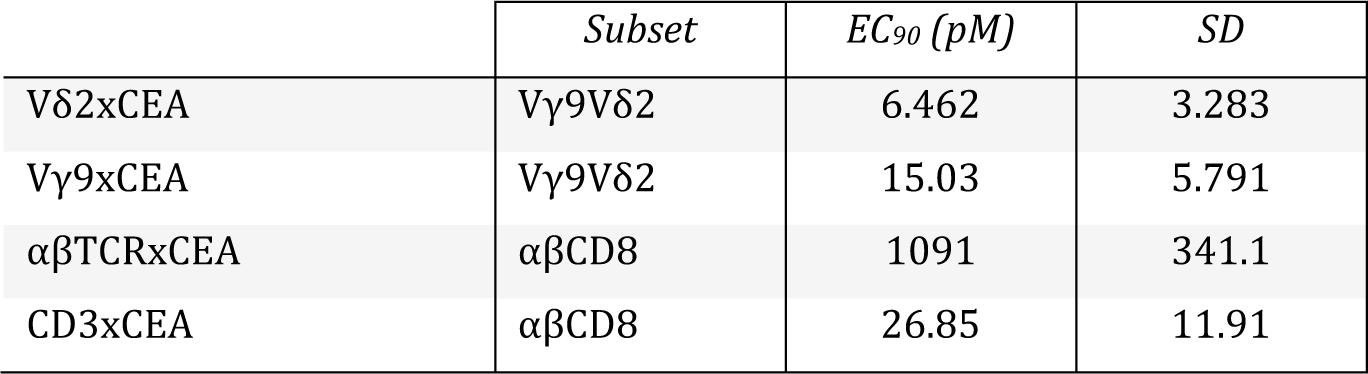
EC_90_ calculated from CD69 expression curves after activation of T-enriched PBMCs by TCEs and SD; n=5 donors

To assess the contribution of the perforin/granzyme killing pathway during TCE engagement, the levels of granzyme B released in the co-culture supernatant at a saturating bsFabs concentration (1 nM) were measured (**Fig. 4E)**. Vγ9-, Vδ2- and αβTCRxCEA bsFabs induced a comparable release of granzyme B by T cell-enriched PBMCs when co-cultured with LS174T cells, which reflects a higher secretion by Vγ9Vδ2 T than αβ T cells as shown by normalizing to the cell number (approximatively 4.7 vs 1.1 fM/cell).

Altogether, these results indicate that Vγ9-, Vδ2- and CD3xCEA bsFabs specifically and efficiently activate the innate-*like* peripheral human Vγ9Vδ2 T cell subset. This activation triggers potent anti-tumor functions, such as perforin/granzyme B-dependent cytolysis with decreased release of pro-inflammatory cytokines.

### Vγ9Vδ2-specific bsFabs fail to induce the expression or release of inhibitory molecules

Finally, the induction of immunosuppressive or pro-tumoral features by bsFabs was investigated. First, the level of PD-1 surface expression was measured in fresh T cell subsets after 24 and 48 h of co-culture with LS174T cells (E/T ratio 10:1) in the presence of a saturating dose of bsFabs (1 nM). Interestingly, γδ and αβ T cells exhibited different responses to bsFabs- induced stimulation. Upregulation of PD-1 expression on Vγ9Vδ2 T cells was observed only with CD3xCEA bsFab at 48 h, but not with Vγ9- or Vδ2xCEA bsFabs. In contrast, αβ T cells displayed a maximum peak of PD-1 expression 24 h after stimulation with both CD3- and αβTCRxCEA (**Fig. 5A**). The supernatants from the co-cultures of fresh T cell-enriched PBMCs and LS174T cells were collected at 24 h and the production of IL-10 and TGF-β1 were measured. At the TCEs EC_90_ (**Table 2**), no IL-10 production was detected in the Vγ9- and Vδ2xCEA bsFabs supernatant activation (**Fig. 5B**).

**Figure 5:**
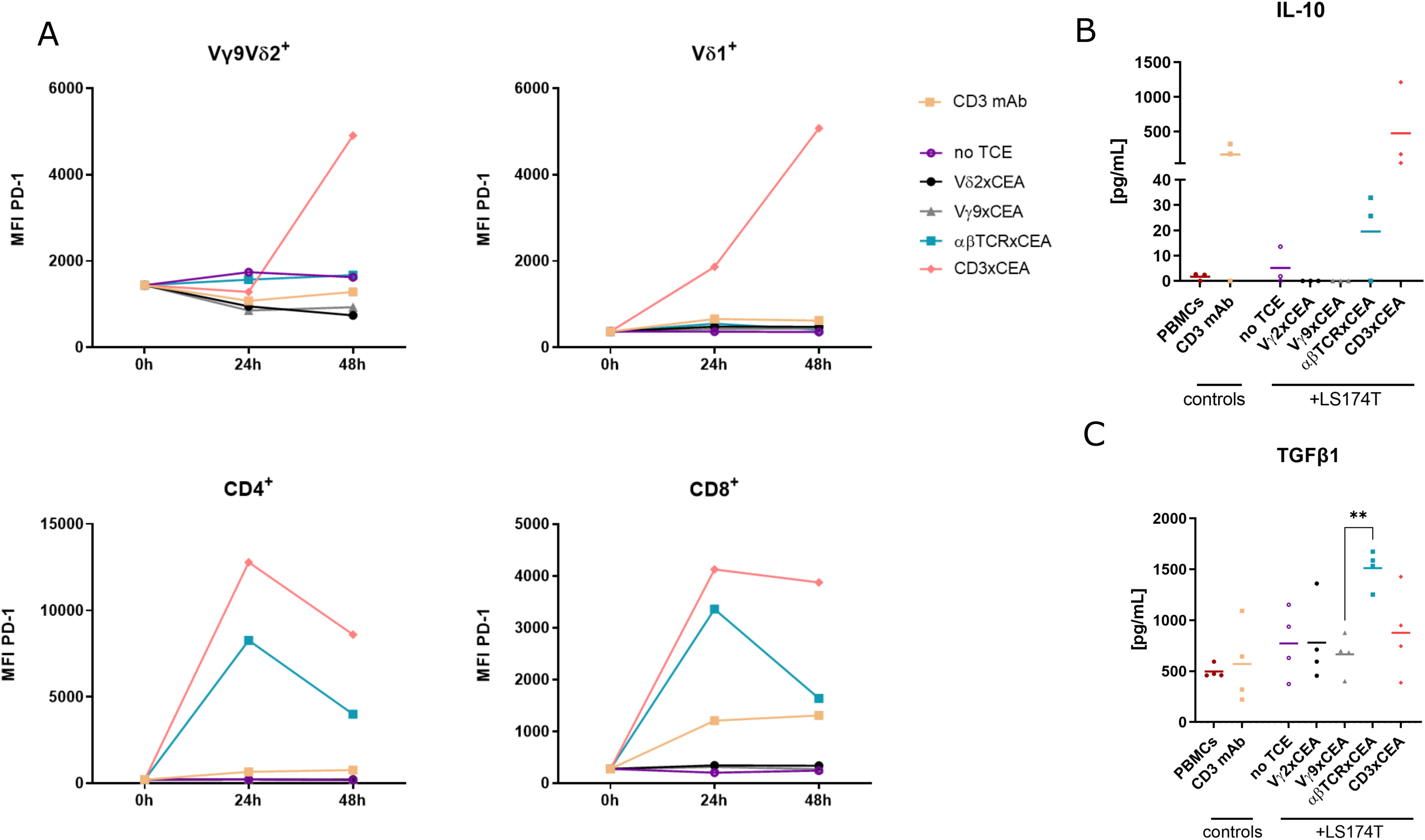
The activation of fresh human peripheral γδ T cells by bsFabs does not induce the expression of exhaustion markers or regulatory molecules. (**A)** Analysis by flow cytometry of surface PD-1 expression (median of fluorescence) on fresh T-cell enriched PBMCs after 0, 24 or 48 h of co-culture with LS174T cells (E/T ratio 10:1) in the presence or absence of xCEA bsFabs (1nM), gated on Vδ2^+^, Vδ1^+^, CD4^+^ or CD8^+^ cells; n=2 donors. Data presented are mean. (**B)** IL-10 concentration measured in the 24 h supernatant of fresh T-cell enriched PBMCs co-cultured with LS174T cells (E/T ratio 10:1), in the presence or absence of xCEA bsFabs at their EC_90_ (calculated from fig.4A); n=3 donors. Data presented are mean. (**C**) Measurement of total TGFβ1 in the 24 h supernatant of fresh T-enriched PBMCs co-cultured with LS174T in the presence or absence of xCEA bsFabs (1nM); n=4 donors. statistical analysis performed using one-way ANOVA tests (ns *P>* 0.05; \**P*< 0.05; **P< 0.01; \*\*\*\**P*< 0.0001). Data presented are mean.

Interestingly, the specific activation of αβT cells by the αβTCRxCEA bsFab showed a distinct pattern with a significant production of TGF-β1, while the activation by the two Vγ9Vδ2 TCR-specific bsFabs did not induce the production of TGF-β1 (**Fig. 5C**). CD3xCEA bsFab induced similar level of TGF-β1 as Vγ9- and Vδ2-CEA bsFab. Collectively, these results suggest that activation of fresh T-cell-enriched PBMCs by Vγ9Vδ2 TCR-specific bsFabs, in contrast to CD3 and αβTCR bsFabs, does not induce surface expression or release of regulatory molecules known for their negative effects on immune responses within the tumor environment.

## Discussion

The ever-increasing number of TCEs progressing to clinical trials and the recent approvals of multiple CD20 and BCMA TCEs for the treatment of hematological malignancies highlight the considerable therapeutic potential of tumor-specific T cell agonism. These immunotherapies suffer however from severe limitations such as side effects or non-optimal efficacy leading the patient to discontinue therapy. Cytokine release syndrome (CRS), infections and neurotoxicity are common side effects which occurrence and severity depend on TCE and indications^39, 40^. Solid cancer treatment by TCEs increases the challenge due to a lower accessibility and a higher risk of off-target toxicity^41, 42^.

This mechanistic study aimed at investigating alternative TCE strategies to reduce side effects while keeping a potent anti-tumor response in solid cancers by specifically targeting human γδ T cell subsets which can account for up to 50% of tissue-resident T cells and for an important proportion of tumor-infiltrated T cells (TILs)^27, 43–46^. We performed an *in vitro* comparative functional study of TCEs targeting all T cells, or either conventional αβ T cells or unconventional Vγ9Vδ2 T cells. The response of these T cell subsets to CD3 TCEs or TCEs specific of each T cell population was investigated; more precisely, their activation level, cytokine production, tumor cell-killing capacity and plasticity to display an immunomodulatory phenotype were compared.

The use of *ex vivo*-expanded T cells, with similar purity ranges, allowed us to study and compare the specific potential of CD3-based TCEs on both αβ T and Vγ9Vδ2 T cells with comparable E:T ratios. Our results showed that the activation of Vγ9Vδ2 T cells by the CD3 bsFab outperformed that of αβ T cells based on a higher surface mobilization of the degranulation marker CD107a and enhanced maximal tumor cell killing. Furthermore, treatment of T cell enriched PBMCs with the CD3 TCE resulted in the activation of all T cell subsets and confirmed the higher sensitivity of Vγ9Vδ2 T cells to CD3 stimulation when compared to αβ T cells. Importantly, the Vγ9Vδ2-specific TCEs were as potent and efficient as the CD3 bsFab to elicit both Vγ9Vδ2 T-cell activation and cytolytic activity, in an antigen-dependent manner. In contrast, the αβ T-cell response to the αβTCR-specific TCE was found to be much less potent than to the CD3 TCE. This could be related to the different affinities of these two TCEs for αβ T cells (≍7 vs 100nM for CD3xCEA vs αβTCRxCEA bsFabs) or to their different epitopes on the full αβTCR, thereby bringing an overall different binding interface known to influence the tightness of the immune synapse and ultimately the TCE potency^47^.

The cytolytic activity of all expanded T cell subsets induced by all bsFabs involved the Perforin/Granzyme pathway as previously reported for other CD3-based TCEs on CD4^+^ and CD8^+^ T cells^4^. We showed that within 30 minutes after addition of any bsFabs, both αβ T and Vγ9Vδ2 T cells have their perforin granules relocated to the immune synapse. The difference of killing activity levels that we measured for the two T cell subsets treated with the CD3 bsFab could be explained by various parameters such as the effector differentiation state and the number of responding T cells, or the intrinsic Granzyme B loading^48^. Circulating Vγ9Vδ2 T cells have an innate-*like* and memory phenotype^49^. Consequently, they require a lower threshold for early antigenic activation and can bypass the need for CD28 costimulation to acquire a potent cytotoxic function as confirmed by our experiments using a monoclonal anti-CD3 mAb. Interestingly, several studies have shown that the lack of a costimulatory signal 2 leads to a non-optimal CD4^+^ / CD8^+^ T cell stimulation by CD3 TCEs and that an additional tumor-specific CD28 stimulation is a promising strategy to improve antitumor activity of CD3 bispecific T cell engagers^50, 51^.

A major adverse effect triggered by TCE-based therapies is the massive production of inflammatory cytokines associated with an over-activation of the immune system, also known as Cytokine Release Syndrome (CRS). Higher grade CRS is most often manageable but requires patient hospitalization and can be life-threatening for the patient^52–54^. Recent engineering strategies aim at decreasing CD3 binding domain affinities or designing conditionally active TCEs^55–59^. An alternative strategy to improve the safety profile of bispecific TCEs while keeping high antitumor activity is to selectively recruit highly cytotoxic T cell subsets such as γδ T cells rather than indiscriminately recruit all T cells (reviewed in ^60, 61^). We compared the effects of specific stimulation of Vγ9Vδ2 T cells by Vg9- or Vd2-based TCE and CD3-based TCE in T cell-enriched PBMCs. While both triggered a similar level of Vγ9Vδ2 T cell activation, specific stimulation resulted in lower levels of a panel of secreted immune effectors such as Granzyme B, IFN-γ and TNF-α, at levels correlated with the low Vγ9Vδ2 : tumor cell ratio. Similarly, the secretion of IL6, one of the most aversive cytokines in cancer immunotherapies^62^, was dramatically decreased when only Vγ9Vδ2 T cells were engaged. In addition, Vγ9- or Vδ2xCEA bsFab treatment blocked the induction of immunomodulatory cytokines (e.g., regulating Treg activation) such as IL-2, IL-10 or TGFβ1 while the CD3 and αβ TCR TCEs induced at least 2 of these 3 cytokines. Finally, the induction of PD-1 expression on T cell surface following activation by bsFabs was found markedly different between the different T cell subsets. αβ T cells rapidly upregulated PD-1 when treated with either CD3 or αβ TCR bsFabs while the γδ T cell response to the CD3 TCE was delayed. More strikingly, PD-1 expression was not induced on Vγ9Vδ2 T cells activated by Vγ9- or Vδ2xCEA bsFabs. This looks particularly encouraging while it has recently be reported that continuous stimulation with TCE can induce CD4^+^/CD8^+^ T cell exhaustion and ultimately results in resistance to therapy^63^.

Altogether, our results indicate that the specific activation of Vγ9Vδ2 T cells by TCEs not only induces a strong cytotoxic activity, but also (i) triggers low production of IFN-γ, TNF-α and IL-6, which should reduce the risk of CRS as compared to large-spectrum TCEs which engage CD3 and subsequently all T cells without discrimination, (ii) is characterized by no to minimal release of modulatory cytokines such as IL-2, IL-10 or TGF-β, and (iii) does not induce the expression of exhaustion markers such as PD-1, suggestive of a durable effective cytotoxic phenotype. This study therefore fully supports the development of novel γδ T bispecific cell engagers, the most advanced of which are currently being evaluated in clinical trials^60, 61^.

As mentioned above, human γδ T cell subsets can account for an important proportion of TILs. While TILs are known to play a key role in effective antitumor immunity, the persistence of an immunosuppressive microenvironment that maintains TILs in an anergic or exhausted state is a major cause of tumor control escape. With this in mind, it would be interesting to generate TCEs that specifically target other human γδ T cell subsets, such as Vδ1^+^ or Vδ3^+^ T cells, which have also been evidenced for their contribution to antitumor responses especially in solid tumors^46, 64^, and study the response of TILs to these TCEs. Would they not be potent enough to overcome the TME immunosuppression impact, combination therapies of γδ T-specific TCEs with allogeneic Vγ9Vδ2 T cell transfer from healthy donors might help enhancing the efficacy of TCEs without additional toxicity^65^.

## Acknowledgements

We thank the Cell and Tissue Imaging Center of the University of Nantes (MicroPICell) for imaging and the Cytocell Cytometry Center of Nantes for technical assistance. We thank Dominique de Chalain and Francis Duffieux (Sanofi) for the purification of the TCEs, Laurent Vidard (Sanofi) for providing *ex-vivo* expanded Vγ9Vδ2 T cells, and Oliver Hijano Cubelos (Sanofi) for the statistical analysis. We thank Margareta Wilhelm for critical reading of the manuscript.

## Authors’ contributions

Conception and design of the project: ES and EV; Analysis and interpretation of the data: LB, ES, EV; Generation and acquisition of data: LB with inputs of E.Se, CB and DB; Study methodology: LB, ES, and EV. Statistical analysis: LB. Writing, reviewing and editing were performed by LB, ES and EV, with inputs from DB and CB.

---

**Suppl Fig 1:**
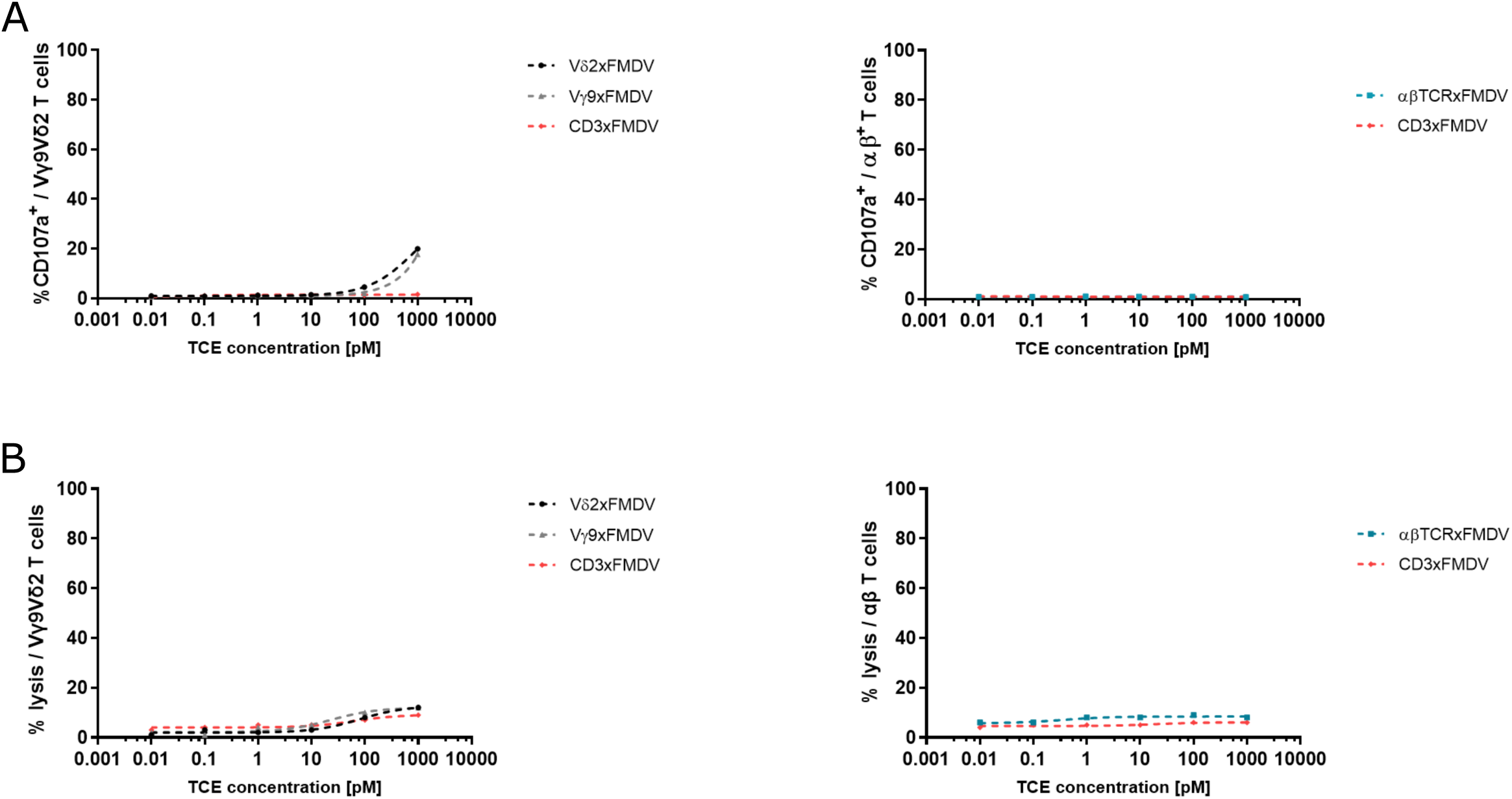
TCE molecule activate *ex vivo* expanded T cells in an antigen-dependent manner. (**A**) Analysis by flow cytometry of CD107a surface expression on *ex vivo*-expanded human Vγ9Vδ2 or αβ T cells after 4 h of co-culture with LS174T cells (E/T ratio 1:1), plus xFMDV TCEs; n=3 unpaired donors. (**B**) LS174T cells were loaded with^51^ Cr and co-cultured for 4 hours with Vγ9Vδ2 T cells or αβ T cells (E/T ratio 10:1), plus xFMDV bsFabs. ^51^Cr release was measured in culture supernatants; n=4 unpaired donors. Data presented from one representative donor.

**Suppl Fig 2:**
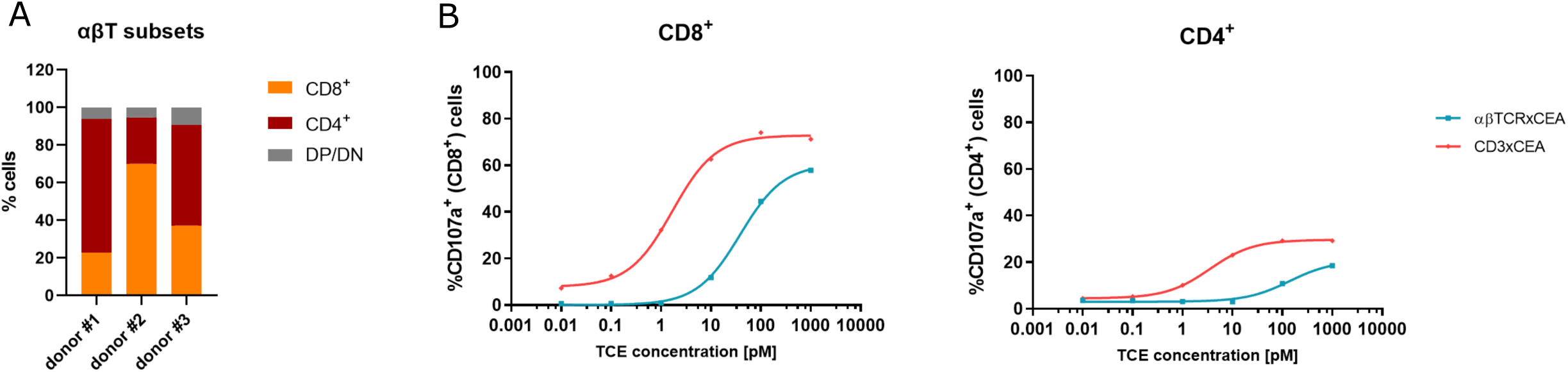
bsFabs mainly activated CD8^+^ among the αβ T cells. (**A**) CD4^+^ and CD8^+^ T cells frequencies in *ex vivo* expanded αβ T cells measured by flow cytometry; n=3 donors. DP/DN = double positive/double negative. (**B**) Flow cytometric analysis of CD107a surface expression on *ex vivo*-expanded CD4^+^ and CD8^+^ T cells after 4 h of co-culture with LS174T cells (E/T ratio 1:1), plus xCEA bsFabs; n=3 donors. Data presented from one representative donor.

